# A fungal powdery mildew pathogen induces extensive local and marginal systemic changes in the *Arabidopsis thaliana* microbiota

**DOI:** 10.1101/2021.02.26.432829

**Authors:** Paloma Durán, Anja Reinstädler, Anna Lisa Rajakrut, Masayoshi Hashimoto, Ruben Garrido-Oter, Paul Schulze-Lefert, Ralph Panstruga

**Affiliations:** Max Planck Institute for Plant Breeding Research, Department of Plant-Microbe Interactions, Carl-von-Linné-Weg 55, 50829 Cologne, Germany; Cluster of Excellence on Plant Sciences, 40225 Düsseldorf, Germany; RWTH Aachen University, Institute for Biology I, Unit of Plant Molecular Cell Biology, Worringerweg 1, 52056 Aachen, Germany; Graduate School of Agricultural and Life Sciences, The University of Tokyo, 113-8657 Tokyo, Japan

**Keywords:** *Arabidopsis thaliana*, *Golovinomyces orontii*, gnotobiotic plant system, microbial multi-kingdom interactions, plant microbiota, powdery mildew, synthetic community

## Abstract

- Powdery mildew is a foliar disease caused by epiphytically growing obligate biotrophic ascomycete fungi. How powdery mildew colonization affects host resident microbial communities locally and systemically remains poorly explored.
- We performed powdery mildew (*Golovinomyces orontii*) infection experiments with *Arabidopsis thaliana* grown in either natural soil or a gnotobiotic system and studied the influence of pathogen invasion into standing natural multi-kingdom or synthetic bacterial communities (SynComs).
- We found that after infection of soil-grown plants, *G. orontii* outcompetes numerous resident leaf-associated fungi. We further detected a significant shift in foliar but not root-associated bacterial communities in this setup. Pre-colonization of germ-free *A. thaliana* leaves with a bacterial leaf-SynCom, followed by *G. orontii* invasion, induced an overall similar shift in the foliar bacterial microbiota and minor changes in the root-associated bacterial assemblage. However, a standing root SynCom in root samples remained robust against foliar infection with *G. orontii*. Although pathogen growth was unaffected by the leaf SynCom, fungal infection caused a more than two-fold increase in leaf bacterial load.
- Our findings indicate that *G. orontii* infection affects mainly microbial communities in local plant tissue, possibly driven by pathogen-induced changes in source-sink relationships and host immune status.

## Introduction

Unlike plants grown under germ-free laboratory conditions, healthy plants in nature live in association and interact with a multitude of microorganisms belonging to several microbial classes, such as bacteria, fungi, oomycetes and protists, collectively called the plant microbiota (Bulgarelli *et al*., 2013). Marker gene amplicon sequencing has served as an important tool for taxonomic profiling and quantitative surveys of microbial assemblages associated with different plant organs over a range of environmental and experimental conditions, revealing community composition and major factors explaining community structure (Hacquard & Schadt, 2015; Thiergart *et al*., 2020). Root-associated bacterial assemblages, assessed by amplicon sequencing of the *16S* rRNA marker gene, are defined by a specific subset of bacteria that originate mainly from the highly diverse soil biota. These communities are characterized by a robust taxonomic pattern at the phylum rank, which comprises the dominant Proteobacteria as well as Actinobacteria, Bacteroidetes, and Firmicutes (Bulgarelli *et al*., 2012; Lundberg *et al*., 2012; Hacquard & Schadt, 2015). The main factors governing differences in root assemblages are, in order of importance, soil type (edaphic factors such as pH), plant species/genotype, and plant age (Lauber *et al*., 2009; Hacquard *et al*., 2015; Müller *et al*., 2016; Finkel *et al*., 2017). For leaf-associated bacterial assemblages, the seeding source is less defined because microbiota members can originate from aerosols, insects, and soil as well as upward microbial migration from the root (Vorholt, 2012; Müller *et al*., 2016). Although root- and leaf-associated microbiota share a similar phylum-level taxonomic composition, the overall community structure in leaves is subject to larger fluctuations. Similar to bacterial communities, plant-associated fungi have been studied at the community level by amplicon sequencing of the internal transcribed spacer (ITS) region between the small- and large-subunit rRNA genes or the *18S* rRNA gene (Bazzicalupo *et al*., 2013). Plant-associated fungal assemblages are dominated by members belonging to the Ascomycota and Basidiomycota phyla, and these communities display more stochastic variation compared with the bacterial microbiota, with biogeography as the strongest explanatory factor (Coleman-Derr *et al*., 2016; Gao *et al*., 2020; Thiergart *et al*., 2020). In fact, taxonomically structured microbial communities with similar taxa on each plant individual have been described until now only for the bacterial root microbiota (Lundberg *et al*., 2012).

Systematic establishment of plant-derived microbial culture collections of the plant microbiota has enabled microbiota reconstitution experiments with germ-free plants and taxonomically representative synthetic communities (SynComs) that can be used to address principles underlying community assembly and proposed microbiota functions under defined laboratory environments. This experimental approach has proven critical to the advancement of microbiota research (Bai *et al*., 2015; Lebeis *et al*., 2015; Durán *et al*., 2018; Zhang *et al*., 2019a). For example, more than 400 leaf- and root-derived bacterial commensals have been isolated from healthy *A. thaliana* grown in natural soil, comprising 35 bacterial families belonging to the aforementioned four phyla (Bai *et al*., 2015). This microbiota culture collection represents the majority of bacterial taxa that are detectable by culture-independent *16S* rRNA gene community profiling in the *A. thaliana* phyllo- and rhizosphere. Microbiota reconstitution experiments using SynComs from this collection revealed that the bacterial root microbiota provides indirect protection to its host against soil-borne and root-associated harmful fungi and that this protection is essential for plant survival (Duran et al., 2018). Similar reconstitution experiments have shown that bacterial root commensals are necessary for iron nutrition of *A. thaliana* in naturally occurring calcareous soils, where poor bioavailability of this soil mineral nutrient limits plant growth (Harbort *et al*., 2020).

Relatively little is known about how the plant microbiota responds to pathogen invasion. The oomycete leaf pathogen *Albugo* has strong effects on epiphytic and endophytic bacterial colonization in *A. thaliana*. Specifically, α-diversity decreased and β-diversity stabilized in the presence of *Albugo* infection in leaves, whereas they otherwise varied between plants (Agler *et al*., 2016). The effect of *Alb. laibachii* on leaf-associated fungal communities were less consistent and not as clear. Upon foliar defense activation by the downy mildew oomycete pathogen *Hyaloperonospora arabidopsidis*, the host *A. thaliana* specifically promotes three bacterial species in the rhizosphere, namely *Xanthomonas, Microbacterium*, and *Stenotrophomonas* sp., respectively (Berendsen *et al*., 2018). Although separately these bacteria did not affect the host significantly, together they induced systemic resistance against downy mildew and promoted growth of the plant.

The analysis of powdery mildew-induced changes in plant leaf microbiota so far rests on field studies with powdery mildew-infected leaf samples (diseased leaves) in comparison to healthy/less infected leaves in Japanese spindle (*Euonymous japonicus*) (Zhang *et al*., 2019b), pumpkin (*Cucurbita moschata*) (Zhang *et al*., 2018) and English oak (*Quercus robur*) (Jakuschkin *et al*., 2016) (reviewed in (Panstruga & Kuhn, 2019)). In the *E. japonicus* study, the authors noticed a reduction in bacterial and fungal diversity, associated with a general decrease in relative abundance (RA) at the genus level (Zhang *et al*., 2019b). Similarly, the richness and diversity of the fungal community was found to be reduced in pumpkin leaves heavily infected by powdery mildew (*Podospharea* sp.) (Zhang *et al*., 2018). Marked changes in the composition of foliar fungal and bacterial communities were also observed in *Erysiphe alphitoides-colonized* oak (Jakuschkin *et al*., 2016). However, all these studies rely on field samples and natural powdery mildew infections in fluctuating conditions, which complicates deconvolution of microbiota changes caused by changes in environmental factors from those driven by pathogen infection.

Here, we examined the effect of controlled powdery mildew (*G. orontii*) infection on the structure of *A. thaliana* leaf and root microbiota in either soil-grown plants or a gnotobiotic plant system pre-treated with defined root- or leaf SynComs. In both settings, we found major powdery mildew-induced shifts in the composition of the local (foliar) assemblages of fungal (natural soil) and bacterial (natural soil and sterile conditions) communities. In the case of the leaf SynCom, this shift was also associated with a marked increase in bacterial load. Apart from these major changes in the phyllosphere, we also observed a minor systemic effect on the structure of the bacterial root microbiota in conjunction with the leaf SynCom and powdery mildew challenge.

## Materials and Methods

### Plant material

*A. thaliana* Col-0 wild-type (Arabidopsis stock centre accession N60000) was used as model plant system. Seeds were surface-sterilized by treating them with 70% for 20 min and drying them under a sterile hood. Subsequently, the seeds were stratified overnight at 4 °C.

### Microbial strains

The A*t*-SPHERE bacterial strains used in this study have been previously reported (Bai *et al*., 2015) and are summarized in Table S1. *G. orontii* (isolate MPIPZ) was used as powdery mildew infection agent. *G. orontii* was regularly propagated every week on 4- to 5-week-old supersuceptible *A. thaliana eds1* plants. Powdery mildew inoculation was conducted by leaf-to-leaf contact of healthy plants with rosette leaves of heavily infected *eds1* plants as reported previously (Acevedo-Garcia *et al*., 2017).

### Natural soil experiment

*A.thaliana* Col-0 seeds were sown into 7×7 greenhouse pots filled with Cologne Agricultural Soil (CAS) batch 12 and grown under short-day greenhouse conditions for 6.5 weeks (12 pots containing 5 seeds each). Then, half of the plants were inoculated with *G. orontii* and all pots were transferred to a growth chamber (day: 21 °C, 10 h light; night: 19 °C; 70% humidity). At 11 days post inoculation (dpi), bulk soil, rhizosphere, root and leaf samples were harvested as previously described (Figure 1A; (Bulgarelli *et al*., 2012; Bai *et al*., 2015)).

**Figure 1.**
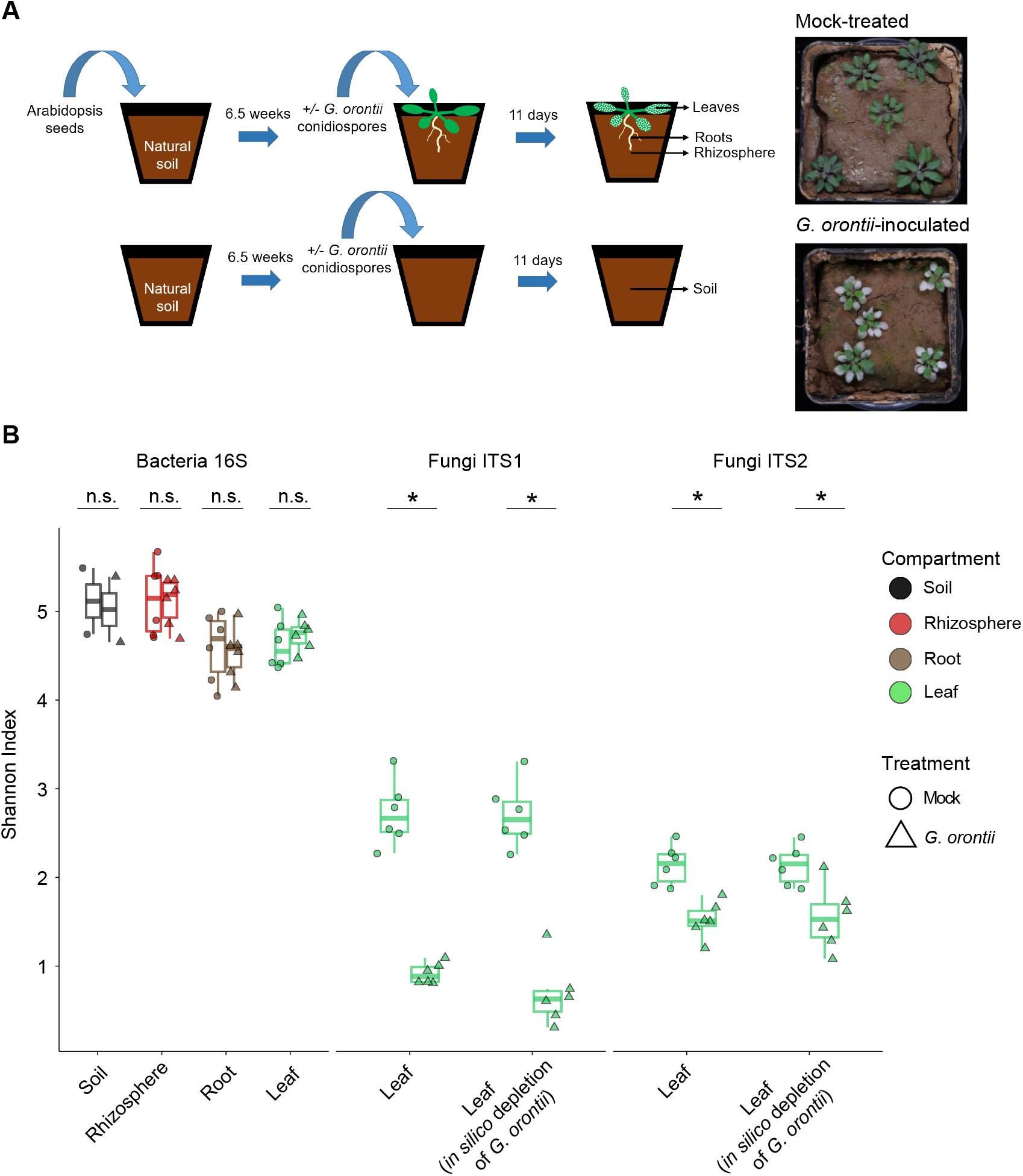
Analysis of α-diversity including all culture-independent samples (ASV-level analysis) for both bacterial and fungal communities. **A)** Schematic experimental set-up and representative pictures of mock-treated and *G. orontii*-inoculated plants. **B)** Boxplots of within-sample diversity (Shannon index) for each compartment and condition. Significant differences in bacterial community diversity within compartments are marked with an asterisk (Student’s *t*-test, * *P*<0.05; n.s., not significant).

### SynCom experiments in gnotobiotic system

Calcined clay was washed several times with tap water followed by MiliQ water. After removal of the liquid, the calcined clay was autoclaved following a liquid cycle (121 °C, 20 min), and oven-dried at 60 °C for 4 weeks. Of the washed, autoclaved and dried calcined clay, 100 g were transferred into a previously sterilized Magenta GA-7 plant culture box (Thermo Fisher Scientific, Schwerte, Germany), sealed and autoclaved again, and dried overnight prior to the experiment.

The bacterial strains used were pre-grown for 7 d in Tryptic Soy Broth 50 % (TSB 50 %, Sigma-Aldrich), and washed with 10 mM MgCl2 (series of centrifugations and removal of supernatant, and final resuspension in MgCl2) to remove any byproducts from the bacterial cultures. A total of 88 and 103 bacterial strains were selected, which were differentiable based on the their *16S* sequence (see Table S1), for the root and leaf SynComs, respectively, and mixed in three separate inputs. For each of the three bacterial mixes, the OD600 was set to 0.5 (root SynCom) and to 0.2 (leaf SynCom) with 10 mM MgCl2. A sample of each of the bacterial inputs was taken as a reference for bacterial community composition.

For the root SynCom, 70 mL ½ MS media (including vitamins without sucrose, pH 7, Sigma-Aldrich GmbH, CatNo M5524-1L, Taufkirchen, Germany) were mixed with 1 mL of the SynCom culture and used to inoculate one calcined-clay-containing Magenta box (n=12). As a control, to a separate batch of magenta boxes only ½ MS media without SynCom culture was added (n=12). Surface-sterilized *A. thaliana* Col-0 seeds were sown at the four corners of the boxes and the lids closed. Plants were grown in a growth cabinet under short-day conditions (day: 22 °C, 10 h light; 70% humidity) for 4.5 weeks. Then, plants were inoculated with *G. orontii* under sterile conditions (n=6, no SynCom culture, and n=6, previously treated with SynCom culture). In addition, some boxes that were either mock- or SynCom-inoculated were opened, but not inoculated with *G. orontii*, serving as a control. At 11 dpi, root and leaf tissues were harvested (Figure 4a).

For the leaf SynCom, 70 mL ½ MS media (including vitamins without sucrose, pH 7, Sigma-Aldrich GmbH, CatNo M5524-1L) were mixed into the calcined-clay-containing magenta boxes, and surface-sterilized *A. thaliana* seeds sown at the four corners of the boxes. Plants were grown in a growth cabinet under short-day conditions (day: 21 °C, 10 h light; night: 19 °C; 70% humidity) for 3.5 weeks. Then, a 10-fold dilution of the leaf SynCom in 10 mM MgCl2 was used to spray five times with a Reagent Sprayer (CAMAG®Glass Reagent Spray, Muttenz, Switzerland) onto the *A. thaliana* plants (see (Bai *et al*., 2015); here: 1 spray volume = approx. 35 μL; n=18). At 14 d after the addition of the SynCom or respective mock treatments, plants were inoculated with *G. orontii* under sterile conditions (n=5, no SynCom, and n=9, previously treated with SynCom culture). In addition, some boxes that were either mock- or SynCom-inoculated were opened, but not inoculated with *G. orontii*, serving as a control. At 7 dpi, root and leaf tissues were harvested (Figure 4a).

### Microbial profiling from plant tissues

Total DNA was extracted from the samples mentioned above using the FastDNA SPIN Kit for Soil (MP Biomedicals, Solon, USA). Samples were homogenized in Lysis Matrix E tubes (MP Biomedicals, Heidelberg, Germany) using the Precellys 24 tissue lyzer (Bertin Technologies, Montigny-le-Bretonneux, France) at 6200 rpm for 30 s. DNA samples were then eluted in 80 μL nuclease-free water and used for bacterial and fungal community profiling (Durán *et al*., 2018). Concentrations of DNA samples were fluorescently quantified, adjusted to 3.5 ng/μL, and samples used as templates in a two-step PCR amplification protocol. In the first step, the V5–V7 region of bacterial *16S* rRNA (primers 799F-1192R), fungal *ITS1* (primers ITS1F-ITS2) and *ITS2* (primers fITS7-ITS4) regions were amplified. Under a sterile hood, each sample was amplified in triplicate in a 25 μL reaction volume containing 2 U DFS-Taq DNA polymerase, 1x incomplete buffer (both Bioron GmbH, Ludwigshafen, Germany), 2 mM MgCl2, 0.3% BSA, 0.2 mM dNTPs (Life Technologies GmbH, Darmstadt, Germany) and 0.3 μM forward and reverse primers. PCR was performed using the same parameters for all primer pairs (94 °C/2 min, 94 °C/30 s, 55 °C/30 s, 72 °C/30 s, 72 °C/10 min for 25 cycles). Afterwards, single-stranded DNA and proteins were digested by adding 1 μL of Antarctic phosphatase, 1 μL xonuclease I and 2.44 μL Antarctic Phosphatase buffer (New England BioLabs GmbH, Frankfurt, Germany) to 20 μL of the pooled PCR product. Samples were incubated at 37 °C for 30 min and subsequently enzymes deactivated at 85 °C for 15 min. Samples were then centrifuged for 10 min at 4000 rpm, and 3 μL of this reaction were used for a second PCR, prepared in the same way as described above, using the same amplification protocol but with the number of cycles reduced to 10, and with primers including barcodes with Illumina adaptors (Table S1). PCR product quality was controlled by loading 5 μL of each reaction on an agarose gel and affirming that no band was detected in the negative control. Afterwards, the replicated reactions were combined and purified as follows: Bacterial amplicons were loaded on a 1.5 % agarose gel and run for 2 h at 80 V; bands with the correct size of ~500 bp were cut out and purified using the QIAquick gel extraction kit (Qiagen GmbH, Hilden, Germany). Fungal amplicons were purified using Agencourt AMPure XP beads (Thermo Fisher Scientific). DNA concentration was again fluorescently determined, and 30 ng DNA of each of the barcoded amplicons were pooled in one library per microbial group. Each library was then purified and re-concentrated twice with Agencourt AMPure XP beads, and 100 ng of each library were pooled together. Paired-end Illumina sequencing was performed in-house using the MiSeq sequencer and custom sequencing primers (Table S1).

### Absolute quantification of microbial load in plant tissues

Genomic DNA from leaves inoculated with SynCom alone (n=9), *G. orontii* alone (n=9), both SynCom and *G. orontii* (n=9), or mock-treated (n=9) was used for absolute quantification of microbial load. Genomic DNA was fluorescently quantified and diluted to an equal concentration of 4 μL. Each sample was subsequently used for absolute quantification *via* PCR (qPCR) by adding SYBR green to monitor the PCR amplification in real time. Each sample was amplified in duplicate: 4 μL of template were mixed with 7.5 μL of SYBR green (brand), together with 1.2 μL of forward primer and 1.2 μL of reverse primer, to a final volume of 15 μL of reaction. For each organism, a specific primer pair was selected: for bacterial assessment, the *16S* rRNA gene (primers 799F-1192R, (Wippel *et al*., 2021)); for *G. orontii*, a GDSL lipase-like gene (primers R263-R264, (Weßling & Panstruga, 2012)); and, for *A. thaliana*, the *At4G26410* gene (primers L658-L659, (Hong *et al*., 2010)). The following program was used for amplification: pre-denaturation for 3 min at 94 °C, followed by 40 cycles of denaturation for 15 s at 95 °C, annealing for 10 s at 62 °C and elongation for 10 s at 72 °C. Melting curve analysis was performed from 55 °C to 95 °C, with a step-wise increase of 0.5 °C. The amount of bacterial and *G. orontii* genes was normalized to the reference plant gene within each individual sample using the 2^-ΔΔCt^ equation (Pfaffl, 2001). Absolute abundances of individual bacteria in SynCom experiments were obtained by multiplying the bacterial gene/plant gene ratio to their individual relative abundances (RAs).

### Processing of *16S* gene and *ITS* region amplicon data

Amplicon sequencing data from the natural soil experiment (plant tissues along with unplanted controls) were demultiplexed according to their barcode sequence using the QIIME pipeline (Caporaso *et al*., 2010). Afterwards, DADA2 (Callahan *et al*., 2016) was used to process the raw sequencing reads of each sample. Unique amplicon sequence variants (ASVs) were then inferred from error-corrected reads, followed by chimera filtering, also using the DADA2 pipeline. Next, ASVs were aligned to the SILVA database (Quast *et al*., 2013) for taxonomic assignment using the naïve Bayesian classifier implemented by DADA2. Next, raw reads were mapped to the inferred ASVs to generate an abundance table, which was subsequently employed for analyses of diversity (using the R package vegan, (Oksanen *et al*., 2007)) and differential abundance using the R package DESeq2 (Love *et al*., 2014).

Sequencing data from SynCom experiments was processed using the Rbec tool (Zhang *et al*., 2021). First, reads were de-replicated into unique tags and subsequently aligned to the reference database, after which initial abundances were assigned to each strain according to the copy number of each exactly aligned tag. Next, tags that were not exactly matched to any sequence in the database were assigned a candidate error-producing reference based on *k*-mer distances. Sequencing reads were then subsampled and an error matrix is calculated using the mapping between subsampled reads and candidate error-producing sequences. The parameters of the error model were recomputed iteratively until the number of re-assignments fell below the set threshold. Strain abundances were then estimated from the number of error-corrected reads mapped to each reference sequence. Next, we generated a count table that was employed for downstream analyses of diversity with the R package vegan (Oksanen *et al*., 2007). Reads assigned to a given strain were normalized by its *16S* copy number. Finally, amplicon data from all experimental systems were visualized using the ggplot2 R package (Wickham, 2016).

## Results

### *G. orontii*-induced changes in root- and leaf-derived natural microbial communities

*A. thaliana* plants (accession Col-0) were grown in Cologne agricultural soil (CAS12; (Bulgarelli *et al*., 2012)) for 6.5 weeks and subsequently rosette leaves inoculated with *G. orontii* conidiospores. Samples of leaf, root, and rhizosphere compartments as well as unplanted soil were harvested at 11 dpi (Figure 1A). Genomic DNA was prepared from these samples and used for PCR-based amplification of the bacterial *16S* rRNA genes and the fungal ribosomal internal transcribed spacer (*ITS*) regions, *ITS1* and *ITS2*. PCR amplicons were subjected to Illumina sequencing and the resulting sequencing data used for the analysis of the bacterial and fungal communities in the respective compartments. Information on the number and RA of amplicon sequence variants (ASVs) in each compartment was used to calculate α-diversity (Shannon index; within-sample diversity), β-diversity (Bray-Curtis dissimilarities; between-sample diversity), ASV enrichment, and taxonomic composition. Consistent with previous reports (Bulgarelli *et al*., 2012; Schlaeppi *et al*., 2014; Thiergart *et al*., 2020), we found the highest bacterial α-diversity in unplanted soil and the rhizosphere compartment, and lower α-diversity in roots and leaves. There was no significant difference in α-diversity between mock-treated and *G. orontii*-inoculated samples for any of the four compartments analyzed (Figure 1B). By contrast, we observed that in leaves, fungal α-diversity, determined *via ITS1* sequences, was significantly (*P*=0.05) decreased in *G. orontii-inoculated* leaves (by ca. 70%; Figure 1B). This drop in α-diversity was retained upon *in silico* depletion of *G. orontii* reads, excluding the possibility that the decrease in species richness was due to an overrepresentation of powdery mildew reads (and thus underrepresentation of reads from other fungal taxa) in the samples. A similar outcome was obtained with *ITS2* amplicons (Figure 1B). Together, these findings suggest that *G. orontii* leaf colonization reduces the diversity of leaf-associated fungal communities whereas α-diversity of bacterial assemblages remains unaltered.

Analysis of β-diversity using principal-coordinates analysis (PCoA) of Bray-Curtis dissimilarities revealed distinctive community compositions in unplanted soil, rhizosphere, root and leaf compartments (30% of variance explained, *P*=0.001, Figure 2A). We noted that β-diversity of leaf-associated bacterial assemblages changes in response to *G. orontii* inoculation (*P*=0.029), whereas bacterial communities remained indistinguishable in the other compartments (Figure 2B and E). This suggests that *G. orontii* exerts a local effect on bacterial community profiles, but not systemically on the bacterial root and/or rhizosphere microbiota. Closer inspection of the *G. orontii*-induced bacterial community shifts in leaves reveals a broad range of bacterial ASVs that show significantly altered RAs (differentially abundant ASVs; *P*<0.05). This shift affects both abundant and low-abundant community members with ASVs that are enriched or depleted (4.4% and 2.5% of differentially abundant ASVs, respectively). The majority of taxa with differential fold changes are either undetectable in the absence of *G. orontii* (enriched taxa) or undetectable following *G. orontii* challenge (depleted taxa; Supplemental Figure 1). Similar to the leaf-associated bacterial commensals, analysis of β-diversity of the leaf-associated fungal community reveals a marked community shift (*ITS1* Figure 2C; *ITS2* Figure 2D). As with the changes in α-diversity (Figure 1B), the differentiation in distinct leaf-associated fungal communities following powdery mildew infection is not due to an overrepresentation of *G. orontii* reads, as indicated by a similar outcome upon *in silico* depletion of the corresponding reads (Figure 2E).

**Figure 2.**
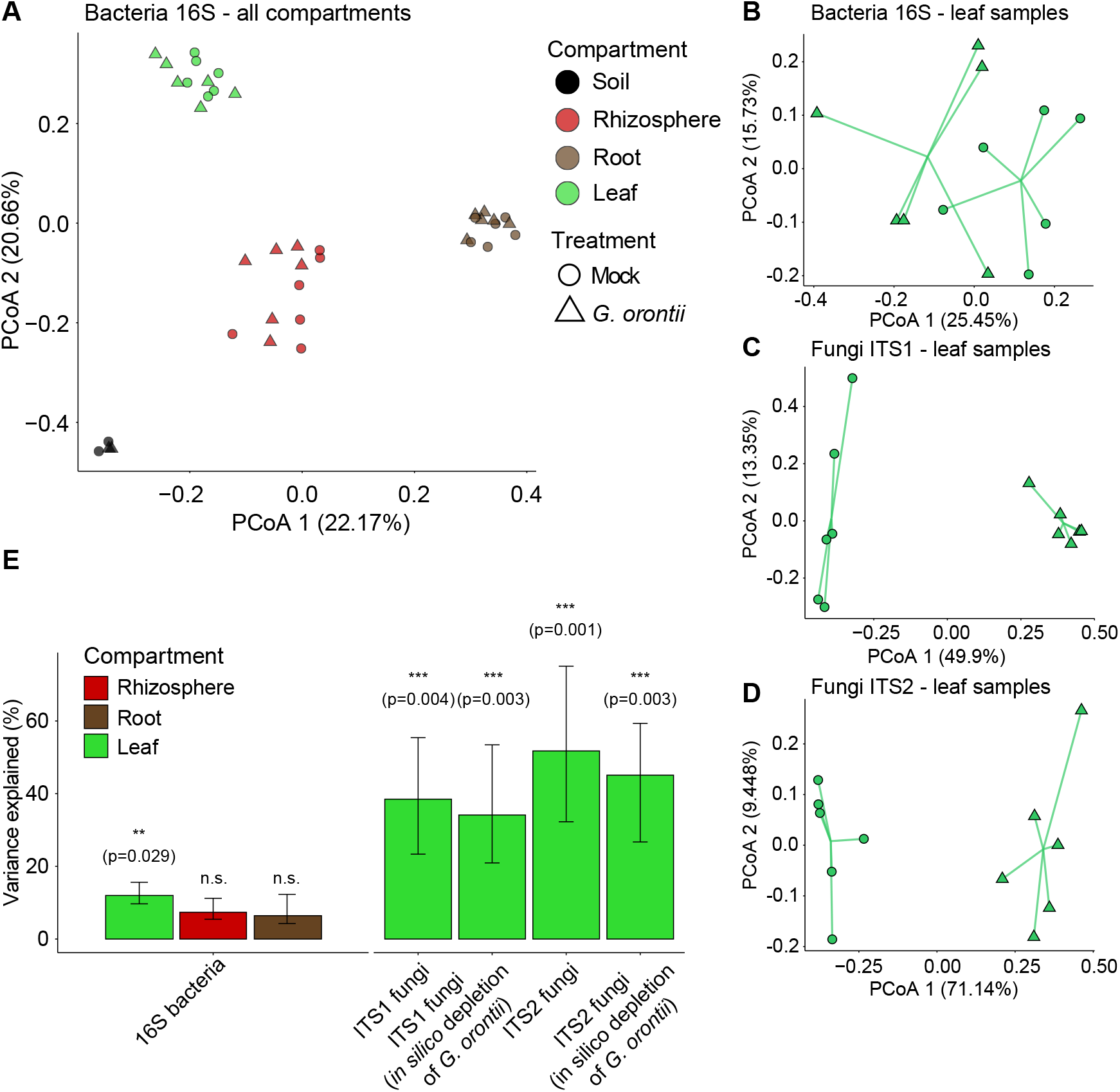
Analysis of β-diversity including all culture-independent samples (ASV-level analysis), for both bacterial and fungal communities. **A)** PCoA plot of Bray-Curtis dissimilarities between bacterial community samples, color-coded by compartment and shaped based on treatment. **B)** Subset of leaf samples where separation between treatments (mock- vs *G. orontii*-treated) can be observed. **C-D)** PCoA plots of fungal communities in leaf samples for *ITS1* (**C**) and *ITS2* (**D**) profiles. **E)** Variance explained of bacterial (left set of columns) and fungal community structure (right set of columns) upon *G. orontii* infection. Note that for fungal communities, an *in silico* depletion of *G. orontii-* assigned reads was performed.

Permutational multivariate analysis of variance revealed that *G. orontii* leaf colonization explained ca. 15% and ca. 34-45% of the variation of the leaf-associated bacterial or fungal communities, respectively (Figure 2E). To explore the changes driving this variation in community structure, we calculated the proportion of bacterial ASVs with differential abundance upon *G. orontii* infection at the order level. For the majority of bacterial orders with multiple differentially abundant ASVs, both enriched and depleted ASVs were found, although we observed more differentially enriched than differentially depleted ASVs. The two orders with the greatest proportion of differentially abundant ASVs were Burkholderiales and Rhizobiales (ca. 33% and 9.5%, respectively; Figure 3A), two abundant bacterial taxa robustly found in *A. thaliana* leaf microbiota (Garrido-Oter *et al*., 2018). We also tested whether differentially enriched or depleted ASVs for a given bacterial order affect its aggregated RA. Whereas the aggregated RA of Burkholderiales, Flavobacteriales and Rhizobiales increases, the aggregated RA of Pseudonocardiales is reduced upon *G. orontii* inoculation (*P*<0.05; Figure 3B). As these four bacterial orders belong to three phyla, Proteobacteria, Actinobacteria and Bacteroidetes, *G. orontii* colonization influences the abundance of phylogenetically distantly related bacterial leaf commensals.

**Figure 3.**
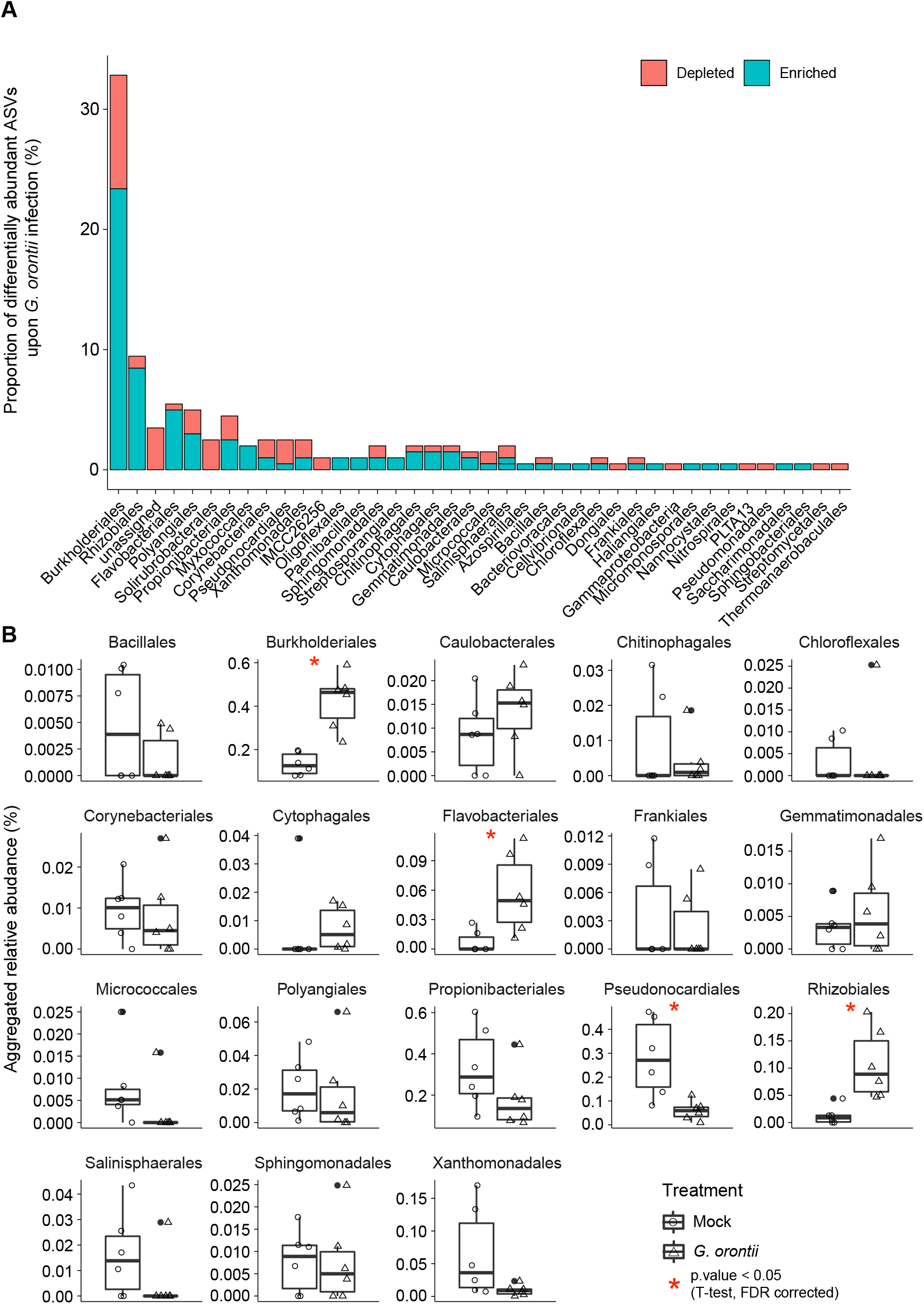
Analysis of bacterial ASVs significantly affected by *G. orontii* infection on leaves. **A)** Proportion of ASVs that significantly changed their relative abundance upon infection with *G. orontii*, grouped by order level, in relation to the overall number of ASVs affected. **B)** Log-transformed aggregated relative abundance of each bacterial order shown in **(A)**, which has both enriched and depleted ASVs, compared between mock and *G. orontii*-treated conditions.

Closer inspection of the *G. orontii*-induced fungal community shift showed that a great proportion (15-34%) of taxa were depleted (Supplemental Figure 2), including abundant and low abundant members of a variety of fungal classes. This result is consistent with the aforementioned *G. orontii*-induced reduction in fungal α-diversity (Figure 1B). However, nine fungal taxa are significantly enriched, with members of the Erysiphales and the Golubeviales showing the highest fold change. Besides *G. orontii*, we noticed the presence of other powdery mildew species (*Erysiphe* spp.) among the enriched fungal taxa. These could either be contaminations in our inoculum or species introduced from the environment in the course of the experiment and “hitchhiking” on the diseased plants.

In summary, we observed a major shift in the resident fungal community in *G. orontii-* infected leaves, characterized by predominantly depleted taxa compared to non-infected plants. In contrast, changes in the leaf-associated bacterial communities were more limited, with both enriched and depleted taxa. No alterations were seen in the systemic root tissue.

### *G. orontii*-induced changes in root- and leaf bacterial SynComs

We examined the impact of *G. orontii* leaf infection using a gnotobiotic *A. thaliana* system and defined *A. thaliana* root and leaf bacterial consortia (root- and leaf-derived A*t*-SPHERE strains), which comprise representatives of the majority of taxa that are detectable by culture-independent *16S* rRNA amplicon sequencing in association with plants grown in natural soil (Bai *et al*., 2015). In this gnotobiotic plant system, *A. thaliana* surface-sterilized seeds were sown on a calcined clay matrix. In the first experiment, we inoculated prior to sowing a defined bacterial consortium consisting of 88 root-derived bacterial commensals (designated here ‘root SynCom’), co-cultivated the consortium with the host for 4.5 weeks, followed by either *G. orontii* conidiospore inoculation or mock treatment. At 11 dpi, leaf, matrix and root samples were harvested and the corresponding DNA preparations subjected to *16S* rRNA amplicon sequencing (Figure 4A, upper panel). PCoA of Bray-Curtis dissimilarities of the bacterial consortia revealed their separation according to compartment (12-22% of variance explained, *P*=0.001, Figure 4B-C). In addition, root-associated consortia were found to be more similar to matrix-associated communities, while still separated from the leaf-resident assemblages (Figure 4B; Supplementary Figure 4). The latter community is likely the result of upward bacterial migration from roots to shoots during co-cultivation (ectopic leaf colonization). In contrast to root samples, leaf samples collected from matrix-inoculated root SynComs differentiated significantly upon *G. orontii* infection (*P*=0.02; Figure 4D and G; Supplementary Figure 4). This pattern is reminiscent of the *G. orontii*-induced impact on *A. thaliana* plants grown in natural soil with an infection-induced change in the bacterial leaf microbiota but non-responsiveness of the root-associated community (Figure 2F).

**Figure 4.**
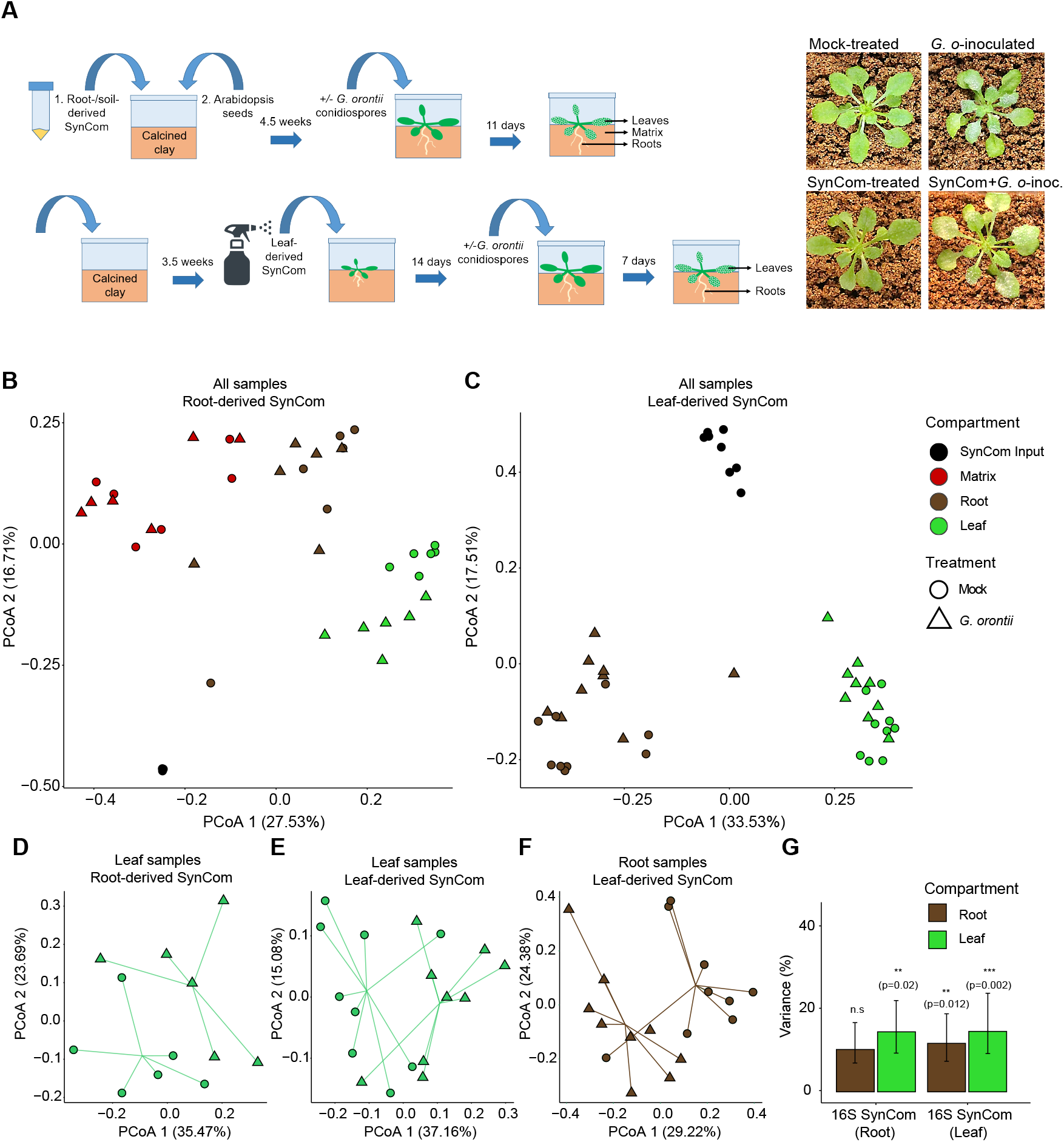
Analysis of beta-diversity including all Root A*t*-SPHERE SynCom samples and all Leaf A*t*-SPHERE SynCom samples based on relative abundances. **A)** Schematic experimental set-up and representative pictures of mock-treated and *G. orontii*-inoculated plants, with and without SynCom. **B)** PCoA of Bray-Curtis distances of all compartment inoculated with Root A*t*-SPHERE strains. **C)** PCoA of Bray-Curtis distances of all compartment inoculated with Leaf A*t*-SPHERE strains. **D-F)** Subset of leaf **(D and E)** and root samples **(F)**, inoculated with either Root A*t*-SPHERE strains **(D)** or Leaf A*t*-SPHERE strains **(E and F)**, showing the effect on bacterial community structure upon *G. orontii* infection (different shapes). **G)** Variance explained on bacterial community structure upon *G. orontii* infection (PERMANOVA analysis on Bray-Curtis dissimilarities).

In the second experiment, *A. thaliana* plants were grown on sterilized calcined clay matrix and at the age of 3.5 weeks we spray-inoculated leaves with a defined bacterial consortium consisting of 103 leaf-derived bacterial commensals (designated here ‘leaf SynCom’). Fourteen days after plant-bacteria co-cultivation, leaves were either inoculated with *G. orontii* conidiospores or mock-treated. At 7 dpi, leaf and root samples were collected and the corresponding microbial DNA preparations subjected to *16S* rRNA gene amplicon sequencing (Figure 4A, lower panel). PCoA of Bray-Curtis dissimilarities of the bacterial consortia shows their separation according to compartments (Figure 4C; Supplementary Figure 5). Closer inspection revealed a significant separation of mock-treated and *G. orontii-* inoculated samples in both roots and leaves (*P*=0.012 and *P*=0.002, respectively; Figure 4E-G; Supplementary Figure 5). The *G. orontii*-induced shift in the leaf-associated bacterial consortium is consistent with the shift of the leaf microbiota observed in *G. orontii-* inoculated plants in natural soil (Figure 2E). However, unlike the absence of a systemic effect on the bacterial root microbiota in natural soil upon leaf powdery mildew infection (Figure 2E), the ectopically located leaf SynCom commensals on roots were responsive to *G. orontii* infection in the gnotobiotic plant system (see Discussion).

To assess potential changes in absolute abundance of leaf-associated bacterial communities following *G. orontii* infection, we performed qPCR of the corresponding leaf samples with PCR primers specific for bacterial *16S* rRNA and a *G. orontii*-specific gene and normalized against an *A. thaliana-specific* amplicon (see Materials and Methods). Unexpectedly, we found an approximately 2.1-fold increase in bacterial load in *G. orontii*-colonized leaf samples, whereas *G. orontii* biomass on leaves in either the presence or absence of the bacterial SynCom remained unaltered (Figure 5A). This was confirmed using PCoA of absolute bacterial abundances in leaf samples spray-inoculated with the bacterial SynCom in the presence or absence of *G. orontii*, explaining 56% of the observed variation (Figure 5B). The increase in bacterial load in *G. orontii*-treated leaves could be seen across multiple taxonomic classes, prominently in Actinobacteria, Alphaproteobacteria and Flavobacteria (Figure 5C, Supplementary Figure 6 for data of individual SynCom strains). We further took advantage of the *G. orontii*-infected plants in the gnotobiotic system to analyze the composition of our inoculum. Although *G. orontii*, as expected, is the dominant taxum of the inoculum, this revealed the presence of three additional fungal genera, including a known hyperparasite of powdery mildews and another powdery mildew species (*Erysiphe* sp., >6% average aggregated RA; Supplementary Figure 7).

**Figure 5.**
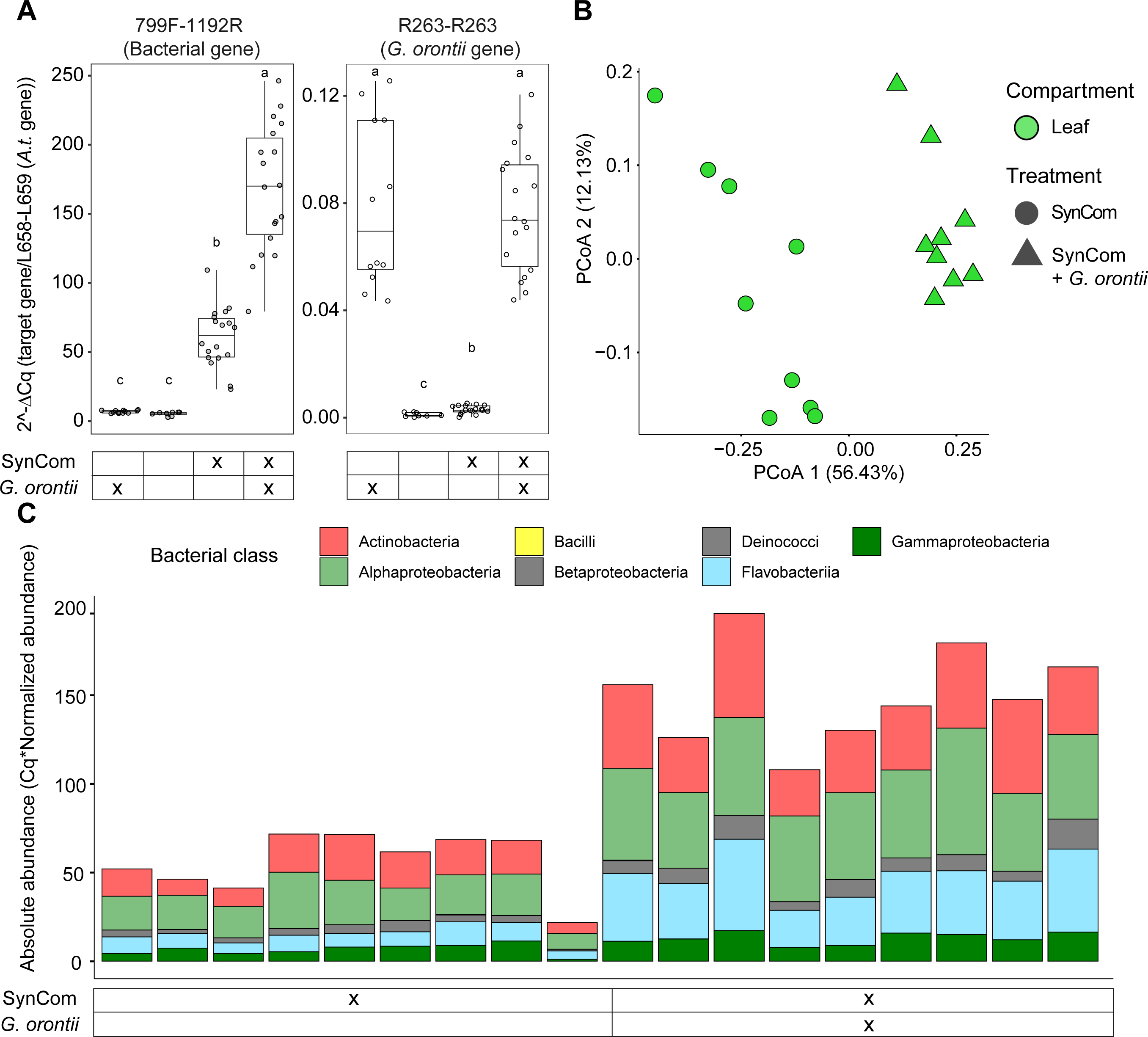
Absolute quantification of bacterial load on leaves inoculated with Leaf A*t*-SPHERE strains. **A)** Bacterial (left panel) and *G. orontii* (right panel) load calculated as the relative quantification of each microbial gene to an *A. thaliana* reference gene. **B)** PCoA plot of absolute abundances of bacterial communities in leaves inoculated with Leaf A*t*-SPHERE strains. **C)** Absolute abundance of each strain utilized in this experiment, color-coded by their taxonomic assignment at the family level, in leaf SynCom-inoculated leaves mock- and *G. orontii*-treated.

## Discussion

In this study, we analyzed in *A. thaliana* the influence of powdery mildew (*G. orontii*) infection on leaf- and root-associated natural and synthetic microbial communities in controlled environments (Figure 1A and 4A). For the experiments conducted in natural soil (Figure 1A), we observed no alteration in bacterial α-diversity in all tested compartments (soil, rhizosphere, root and leaf) following powdery mildew challenge, while α-diversity of the foliar fungal community was strongly reduced (Figure 1B). The more pronounced effect seen with the *ITS1* primers compared to the *ITS2* primer pair was probably a consequence of a greater molecular diversity recovered with the former primers (Bazzicalupo et al., 2013). The substantial reduction in α-diversity of the leaf-associated fungal community upon powdery mildew infection is an indication that the invasive pathogen outcompetes many resident fungal leaf endophytes (15-31% of ASVs), e.g. due to a powdery mildew pathogen-induced shift in sink-source relationships in infected compared to pathogen-free leaves. During a compatible powdery mildew interaction, photosynthetic activity of the plant host is progressively reduced both in cells directly below fungal colonies and in adjacent cells, and this process is associated with an increase in apoplastic invertase activity and an accumulation of hexoses thought to favour pathogen nutrition (Wright et al., 1995; Swarbrick et al., 2006; Eichmann & Hückelhoven, 2008). Alternative explanations may include fungus-specific antibiosis by *G. orontii* or activation of plant immune responses by the fungal invader. The former explanation appears unlikely because the obligate biotrophic pathogen has an unusually low genomic capacity for the biosynthesis of specialized metabolites (Spanu et al., 2010). *G. orontii* pathogenesis stimulates in leaves the accumulation of the defense hormone salicylic acid (SA) at later stages of infection (four dpi) and SA-dependent defense signaling limits hyphal growth and pathogen reproduction (Dewdney *et al*., 2000; Stein *et al*., 2008; Poraty-Gavra *et al*., 2013). For this reason, we consider it plausible that the dramatic reduction in α-diversity of the resident community of asymptomatic leaf-associated fungi in response to *G. orontii* invasion is linked to pathogen-induced and SA-dependent immune responses and/or change in metabolic sink-source relationships. We can, however, not rule out that powdery mildew pathogens deploy secreted effector proteins to antagonize resident endophytic fungi that compete for the same ecological niche (Snelders *et al*., 2018). Potentially different metabolic demands of leaf-associated commensal bacteria compared to fungi together with the ASV-level compensatory changes seen within numerous bacterial orders of the leaf microbiota could explain why the *G. orontii*-induced shift in nutrient availability and immune status does not affect bacterial commensal diversity in the same way.

Consistent with a strong decrease in fungal species richness on *G. orontii*-infected leaves of plants grown in natural soil, we found that the altered fungal community profile is characterized by a reduction in RAs for most and an increase in RAs of a few taxa, while the local bacterial community profile was only slightly affected (Figure 2; Supplementary Figure 3). Leaf-associated fungal endophytes whose RA increases upon *G. orontii* infection comprise mainly one other powdery mildew species (*Erysiphe* sp.) or fungi typically associated with them, such as *Golubevia* sp., which are basidiomycete hyperparasites of powdery mildew pathogens (Russ *et al*., 2021).

Closer inspection of the *G. orontii*-induced shift of bacterial community diversity in leaves revealed 40 bacterial orders, containing several ASVs whose RA is depleted or enriched (Figure 3A and Supplemental Figure 1). However, significant alterations in aggregated ASV-level RAs were seen only in four bacterial orders, with three of them showing an increase and one a decrease (Burkholderiales, Flavobacteriales, Rhizobiales and Pseudonocardiales, respectively; Figure 3B). This pattern suggests widespread compensatory changes at the ASV level within bacterial orders that may allow maintenance of the higher taxonomic structure of the bacterial leaf microbiota during powdery mildew pathogenesis. The fungal pathogen thus mediates local community shifts in the aggregated RAs limited to four bacterial orders belonging to the three main phyla of the *A. thaliana* microbiota (Actinobacteria, Proteobacteria and Bacteriodetes (Bulgarelli *et al*., 2012)). In addition, in our SynCom experiments, colonization by the biotrophic pathogen unexpectedly increased the load of bacterial commensals in leaves approximately 2.1-fold, and this increase was seen roughly equally proportional for all bacterial classes analyzed (Figure 5). The rise in bacterial numbers could be an indirect consequence of the altered sink-source relationships upon infection with the biotrophic pathogen, which turns infected leaves into a metabolic sink associated with an altered availability of non-structural carbohydrates (Wright *et al*., 1995; Swarbrick *et al*., 2006). Consequently, the bacterial commensals might benefit from an accumulation and/or alteration in fluxes of apoplastic hexose sugars and reduction in export of sucrose from the leaf.

In the case of bacteria, changes were only detectable locally (in the infected leaves) but not systemically (in roots or rhizosphere compartments) and only applied to community diversity, while species richness remains unaltered (Figure 1B and 2). The subtle shift in local bacterial diversity could be an indirect consequence of the significant alterations in fungal community richness and diversity upon *G. orontii* invasion. However, at the bacterial order level, we noted a striking overall similarity of the *G. orontii*-induced local bacterial community shifts between the natural soil and bacterial SynCom experiments (significant increases in the RA of Burkholderiales, Flavobacteriales and Rhizobiales and a similar trend for Caulobacterales; Supplementary Fig. 6B). This suggests that our gnotobiotic plant system recapitulates features of the powdery mildew pathogen-induced bacterial community shifts seen in plants grown in natural soil. It further rules out the possibility that these shifts are the result of bacterial immigration into vacated leaf niches that had to be abandoned by the resident fungal endophytes in the course of *G. orontii* pathogenesis.

The only systemic effect on root-associated microbes upon powdery mildew infection reported so far involves nitrogen-fixing rhizobia, which engage in root symbiosis with legumes. In pea (*Pisum sativum*), powdery mildew (*Erysiphe pisi*) colonization was found to result in both a reduction of nodulation and reduced size of root nodules in the leaf-infected plants (Singh & Mishra, 1992). The absence of a systemic effect on the bacterial root microbiota following *G. orontii* challenge differs from *A. thaliana* plants infected with the foliar downy mildew pathogen, the obligate biotrophic oomycete *H. arabidopsidis*. Colonization of leaves with *H. arabidopsidis* promotes the specific enrichment of three bacterial taxa (*Xanthomonas, Stenotrophomonas* and *Microbacterium* spp.) from soil to the rhizosphere, where these three taxa synergistically induced systemic defence responses and promoted plant growth, resulting in enhanced disease resistance against downy mildew (Berendsen *et al*., 2018). Although *G. orontii* and *H. aribidopsidis* are both obligate biotrophic foliar pathogens, only the latter pathogen can infect leaf mesophyll cells and ramify inside this organ, and this may be one reason why only *H. arabidopsidis* can systemically induce changes in the bacterial root microbiota.

Our experiments using germ-free plants, pre-inoculated with a root- or a leaf-SynCom and either mock-treated or challenged with *G. orontii* (Figure 4), recapitulated in parts the pathogen-induced shift in bacterial community profiles of plants grown in natural soil. However, in these experiments also the root samples showed a significant difference in β-diversity of the bacterial root microbiota following *G. orontii* inoculation (Figure 4F and G). As roots were not pre-inoculated in this experiment, the bacteria recovered from root samples must have originated from the leaf inoculation and either reached the below-ground roots in calcined clay accidentally (e.g. by wash-off) or by downward migration, e.g. *via* the plant vasculature. Irrespective of the mechanism(s) underlying ectopic root colonization of the leaf-inoculated bacterial SynCom, this result indicates a (subtle) systemic effect of foliar inoculation with the powdery mildew pathogen on the root-associated bacterial community, which was not seen in the experiment with plants grown in natural soil (see above and Figure 2B). This difference is probably the result of different conditions in the two experimental setups. The SynCom experiment involves a reduced complexity of the bacterial community, allowing us to track the RA of many members with strain-specific resolution rather than the RA of ASVs, which could represent an average of multiple genetically polymorphic strains that share an identical *16S* rRNA gene. Therefore, the higher complexity of the root microbiota in natural soil may mask putative systemic effects on bacterial assemblages induced by *G. orontii* colonization.

It is surprising that the pre-inoculated leaf SynCom does not mediate detectable protective activity against the fungal powdery mildew pathogen, e.g. through activation of defence-associated gene expression in leaves or antagonistic bacteria-fungus interactions (Figure 5A; (Vogel *et al*., 2016; Durán *et al*., 2018)). In roots, the presence of the bacterial microbiota is essential for indirect protection and survival of *A. thaliana* against the otherwise detrimental activity of diverse fungal root endophytes (Durán *et al*., 2018). In *A. thaliana* leaves, several members of the genus *Sphingomonas*, originally isolated from plants, conferred plant protection against the foliar bacterial pathogen *P. syringae* DC3000 and *Xanthomonas campestris* in a gnotobiotic system, whereas no protection was observed by colonization with members of the genus *Methylobacterium* (Innerebner *et al*., 2011). Thus, our results suggest that the bulk of the resident bacterial leaf microbiota of *A. thaliana* does not have a dedicated role in indirect protection against epiphytic *G. orontii* invasion in the tested gnotobiotic system. Whether this is also true for fungal leaf pathogens that colonize mesophyll cells in the leaf interior remains to be tested.

Part of the powdery mildew-induced changes in the *A. thaliana* microbiota could be due to the suppression of plant immunity and reprogramming of host cells for parasitism by the obligate biotrophic pathogen (Schulze-Lefert & Panstruga, 2003). The effect of defence suppression in colonized and neighboring leaf epidermal cells is for example evident by the phenomenon of “induced accessibility”, which refers to enabled host cell entry of non-adapted powdery mildews in the vicinity of established powdery mildew infection sites (Yamaoka *et al*., 1994; Lyngkjær & Carver, 1999b; Lyngkjær & Carver, 1999a; Lyngkjær *et al*., 2001). Powdery mildew-induced alterations of the host transcriptome likely represent the net outcome of counteracting activities of the plant immune system and the fungal intruder (Fabro *et al*., 2008). Although leaf-associated bacterial commensals also extensively reprogram host transcriptomes, stimulating and/or repressing the activity of gene clusters enriched in immunity-and metabolism-associated functions in leaves (Vogel *et al*., 2016), the additional pathogen-specific changes in the host transcriptional profile during infection likely contribute to modulating the quantity and composition of the resident microbiota.

Based on the analysis of our *G. orontii* inoculum on gnotobiotic plants, we noted that approximately 50% of the reads originate from different fungi, including a powdery mildew hyperparasite and another powdery mildew species (*Erysiphe* sp.; Supplementary Figure 7). Our finding highlights the difficulties of maintaining a plant pathogen with an obligate biotrophic lifestyle that must be propagated on living host plants in pure culture. The co-occurrence with these microbes may balance powdery mildew proliferation. These observations might have implications not only for the interpretation of aspects of the data obtained in this work, but also for studies that use other obligate biotrophic plant pathogens.

## Abbreviations

ASV: Amplificon sequence variant
ITS: Internal transcribed spacer
PCoA: Principal coordinates analysis
RA: Relative abundance
SynCom: Synthetic community

## Author contributions

PSL and RP conceived the project. RP, PSL and RGO designed the experiments. ALR, AR and HM performed the experiments with natural soil. ALR, AR and PD performed the SynCom experiments. PD and RGO analyzed the data. PD and RGO created the Figures. RP and PSL wrote the manuscript with support from AR, PD and RGO.

## Data availability statement

Raw demultiplexed sequencing data and corresponding mapping files will be available at ENA accession number PRJEB43139.

---

**Supplementary Figure 1: Changes in RA of bacterial ASVs in *A. thaliana* leaves upon *G. orontii* inoculation.** Each bacterial ASV that showed significant enrichment in leaf samples treated with *G. orontii*, compared to mock-treated samples (*P*<0.05, DESeq package in R) is shown. Each row represents a different ASV, where the size of each dot corresponds to the log-transformed RA in a given sample (columns). The dots are color-coded based on the ASV taxonomic assignment at the class level. The right panel depicts the fold change of each significantly enriched ASV in *G. orontii*-treated leaves, compared to mock control.

**Supplementary Figure 2: Changes in RA of fungal ASVs in *A. thaliana* leaves upon *G. orontii* inoculation, for the ITS1 community profiles.** Each fungal ASV that showed significant enrichment in leaf samples treated with *G. orontii*, compared to mock-treated samples (*P*<0.05, DESeq package in R) is shown here. Each row represents a different ASV, where the size of each dot corresponds to the log-transformed RA in a given sample (columns). The dots are color-coded based on the ASV taxonomic assignment at the class level. The right panel depicts the fold change of each significantly enriched ASV in *G. orontii*-treated leaves, compared to mock control.

**Supplementary Figure 3: Changes in RA of fungal ASVs in *A. thaliana* leaves upon *G. orontii* inoculation, for the ITS2 community profiles.** Each fungal ASV that showed significant enrichment in leaf samples treated with *G. orontii*, compared to mock-treated samples (*P*<0.05, DESeq package in R) is shown here. Each row represents a different ASV, where the size of each dot corresponds to the log-transformed RA in a given sample (columns). The dots are color-coded based on the ASV taxonomic assignment at the class level. The right panel depicts the fold change of each significantly enriched ASV in *G. orontii*-treated leaves, compared to mock control.

**Supplementary Figure 4: Heatmap of bacterial strain-specific RA including SynCom samples (Root A*t*-SPHERE strains).** Warm colors correspond to abundant strains detected in at least one input sample (>0.1% RA). Samples (columns) are grouped by compartment and *G. orontii* treatment.

**Supplementary Figure 5: Heatmap of bacterial strain-specific RAs including SynCom samples (Leaf A*t*-SPHERE strains).** Warm colors correspond to abundant strains detected in at least one input sample (>0.1% RA). Samples (columns) are grouped by compartment and *G. orontii* treatment.

**Supplementary Figure 6: Comparison of absolute abundances of individual Leaf A*t*-SPHERE bacteria to RAs in the culture-independent approach. A)** Absolute abundance (shown as the product of the bacterial/plant gene ratio multiplied with the bacterial RA of each bacterial strain from the Leaf A*t*-SPHERE inoculated in *A. thaliana* leaves without and with *G. orontii*. Boxplots are color-coded based on the taxonomic assignment of each strain at the class level. Significant differences in absolute abundances upon *G. orontii* inoculation are depicted with a red asterisk (Student’s *t*-test, *P*< 0.05, FDR corrected). **B)** Aggregated relative (culture-independent approach) and absolute (culture-dependent approach) abundances of bacterial orders shared between the two experimental set-ups. Significant differences in abundances upon *G. orontii* inoculation are depicted with a red asterisk (Student’s *t*-test, *P*< 0.05, FDR corrected).

**Supplementary Figure 7: Profiling of *G. orontii* inoculum. A)** Number of reads assigned to either bacterial or fungal ASVs in *A. thaliana* leaves inoculated with *G. orontii* **B)** RAs of bacterial ASVs color-coded based on their taxonomic assignment at the class level. **C)** RAs of fungal ASVs from the ITS1 profiles, color-coded based on their taxonomic assignment at the genus level. **D)** RAs of fungal ASVs from the ITS2 profiles, color-coded based on their taxonomic assignment at the genus level.

## Supplementary Table 1

- List of Root A*t*-SPHERE strains used in this study and their taxonomic assignment
- List of Leaf A*t*-SPHERE strains used in this study and their taxonomic assignment
- Primers used in library preparation for amplicon sequencing
- Sequencing primers used for MiSeq sequencing
- Natural soil experiment mapping file for the analysis of community profiles
- SynCom experiments mapping file for the analysis of community profiles
- Primers used in qPCR for microbial absolute quantification in plant tissues

