## Supplementary figures and images for "A fungal powdery mildew pathogen induces extensive local and marginal systemic changes in the *Arabidopsis thaliana* microbiota"

### Supplementary Figures

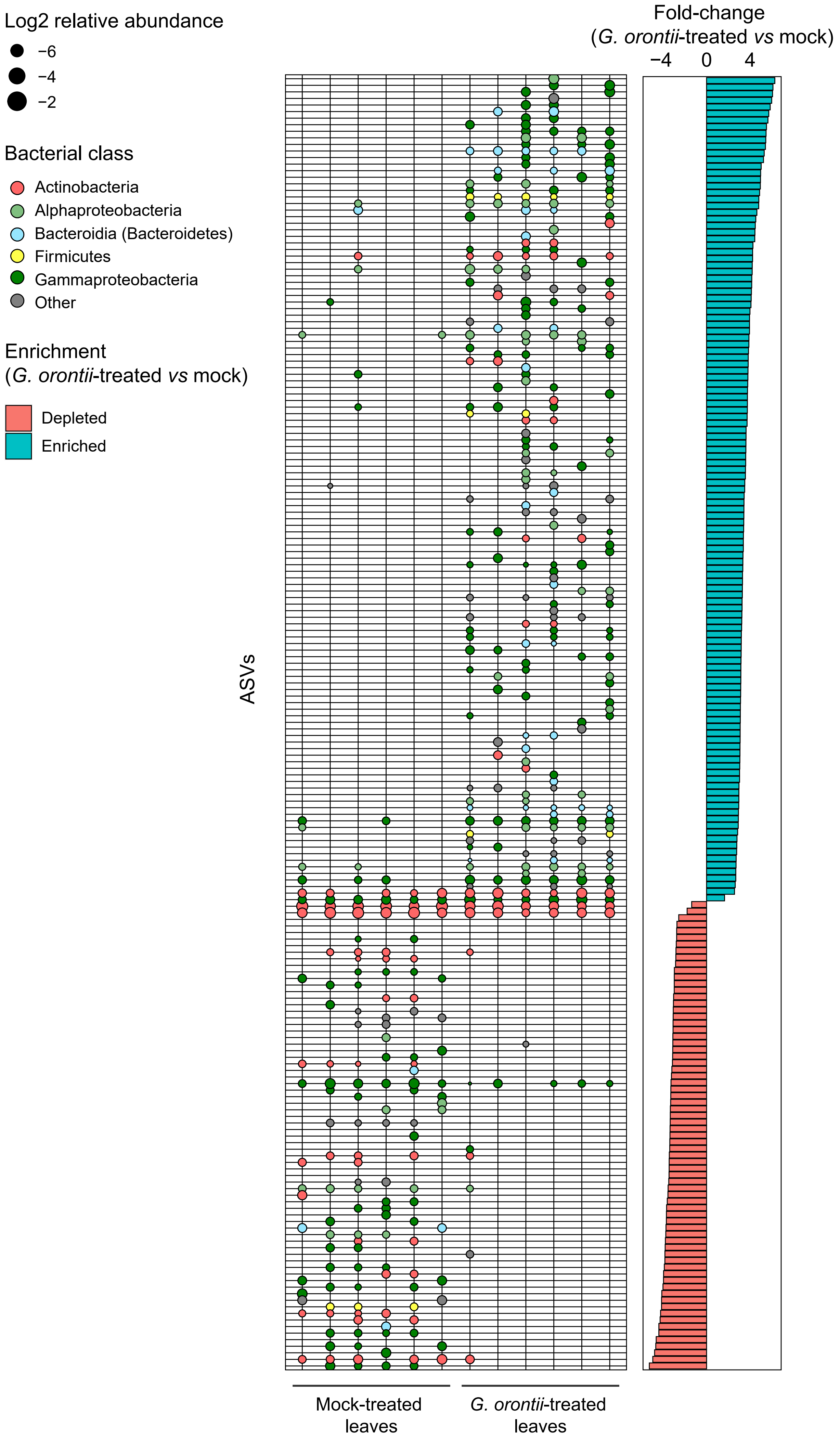

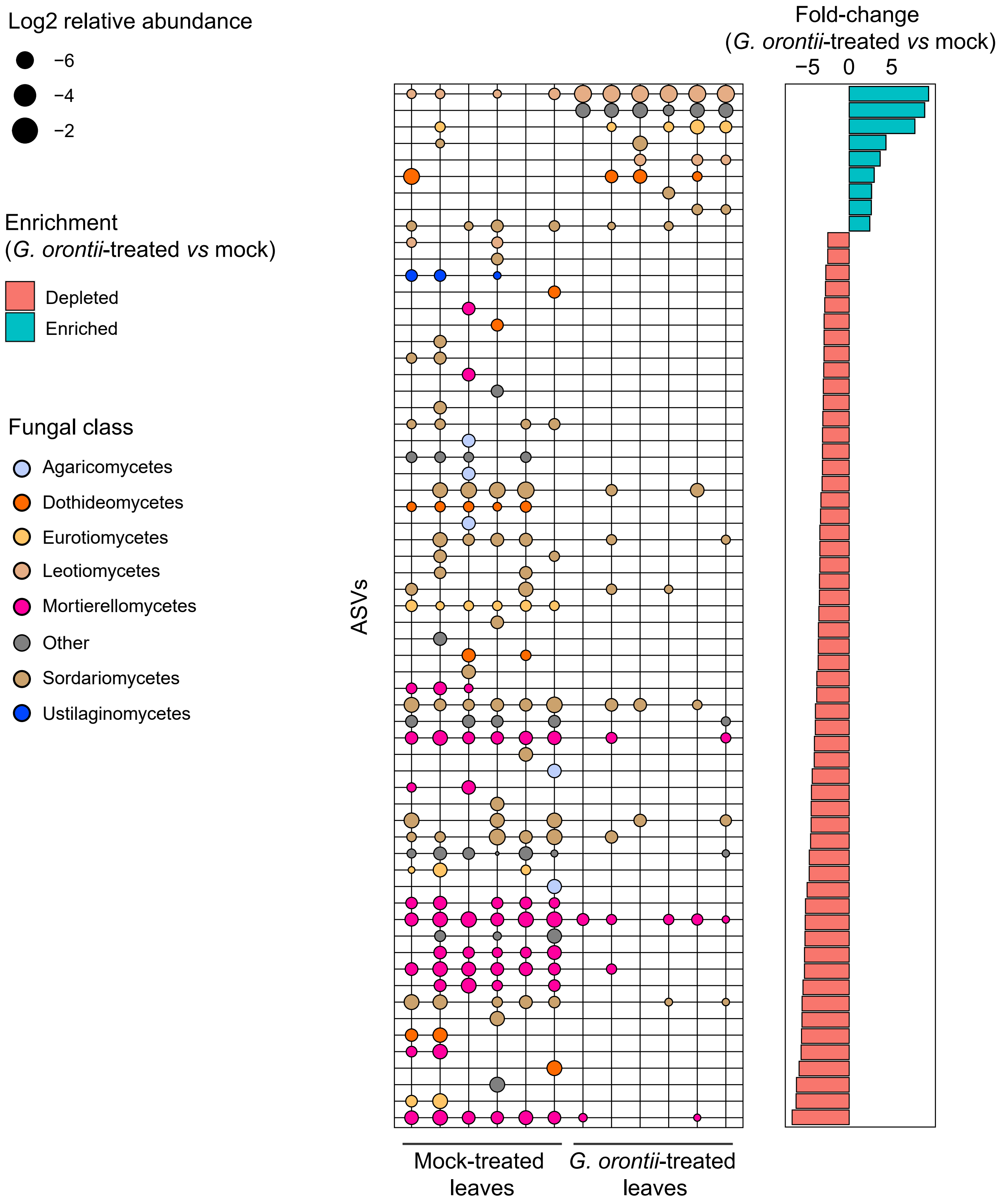

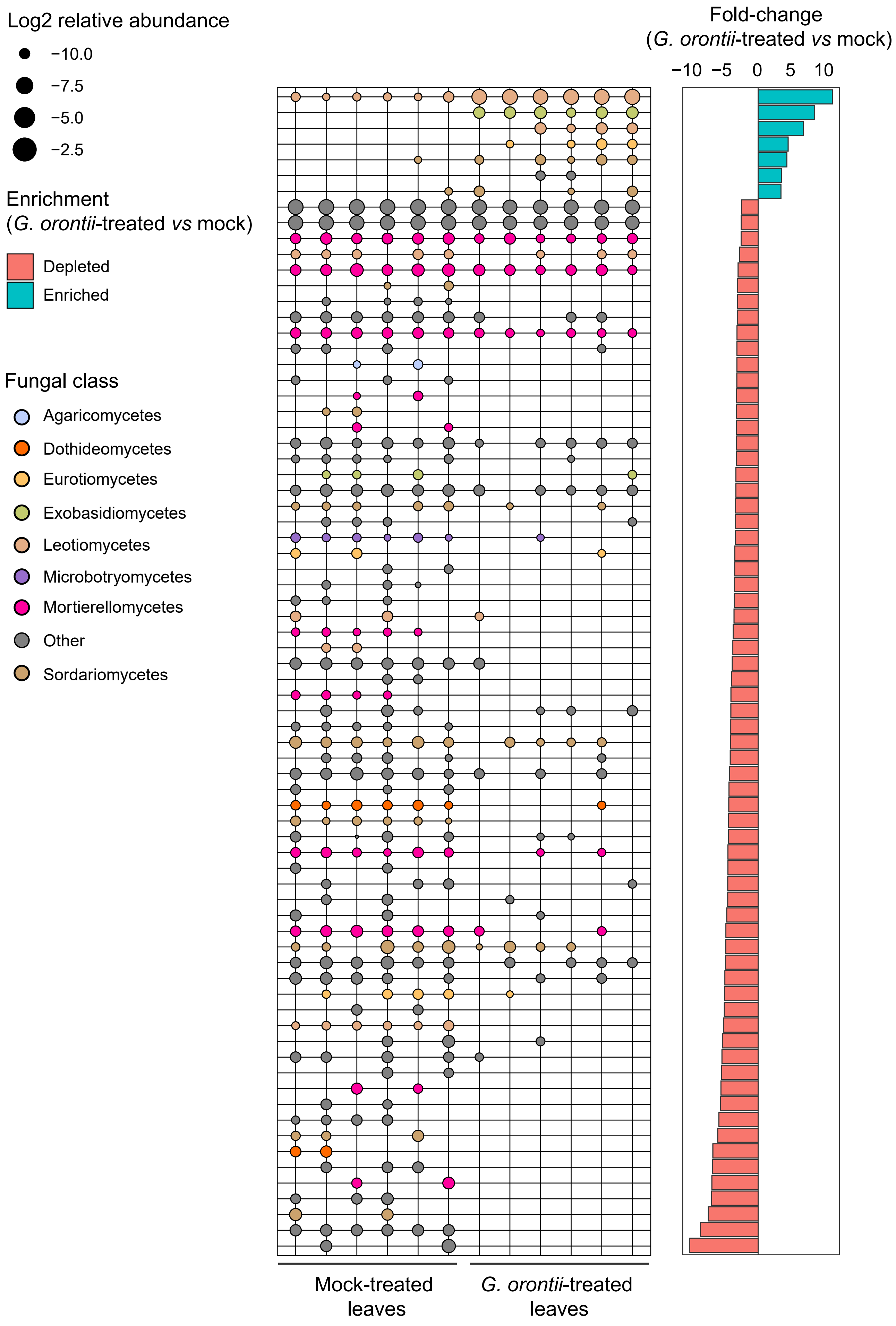

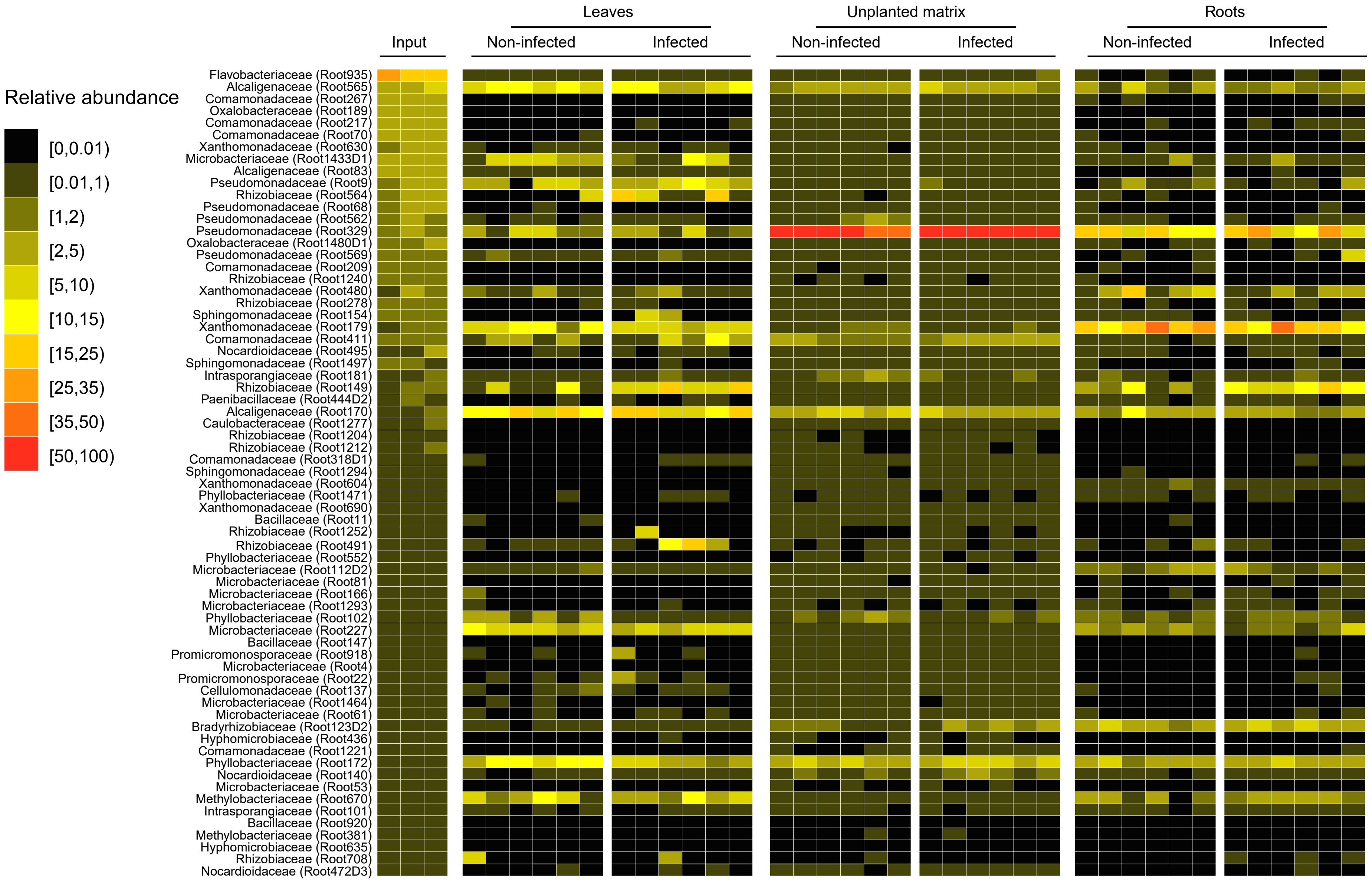

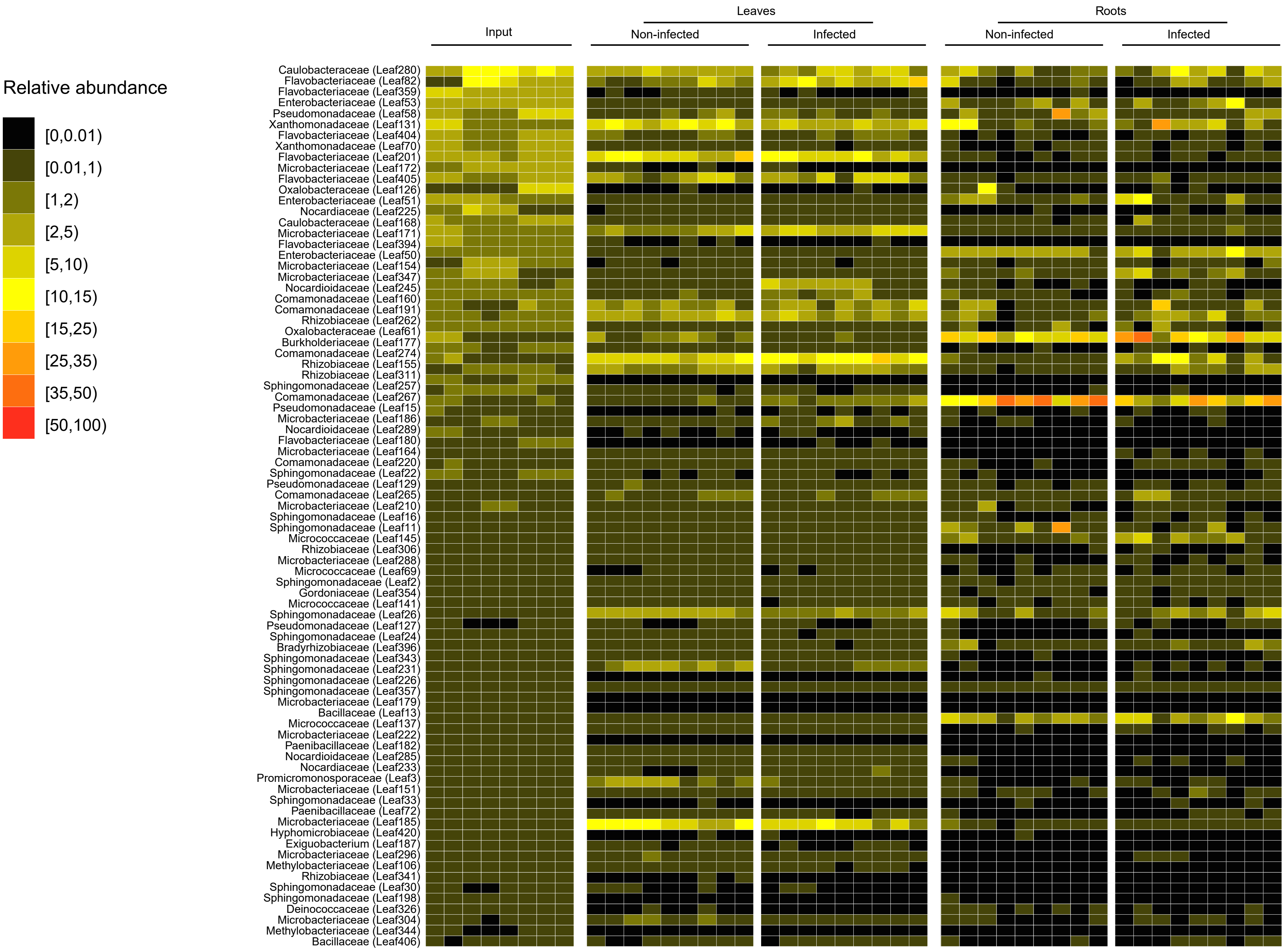

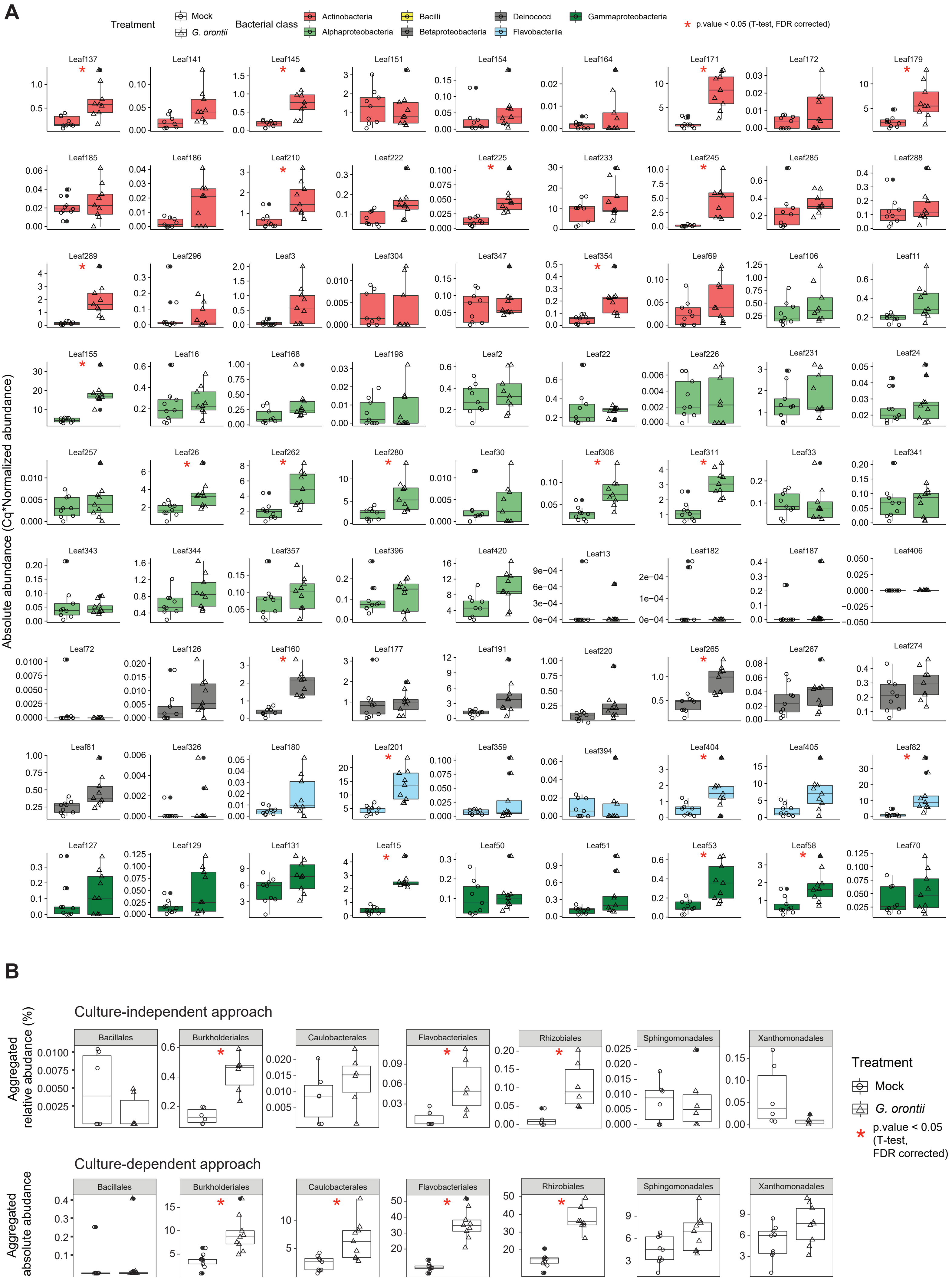

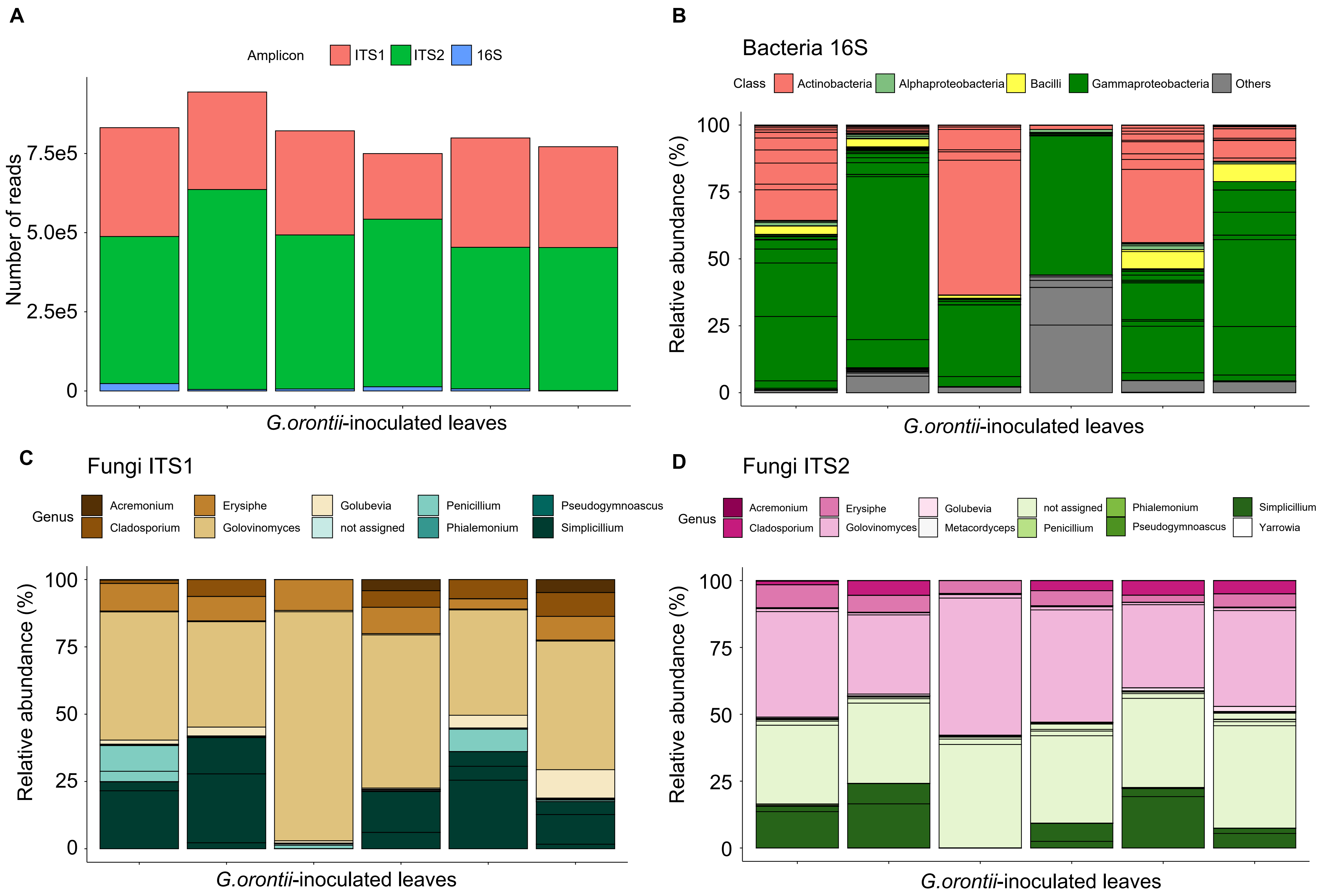
